# Hundreds of viral families in the healthy infant gut

**DOI:** 10.1101/2021.07.02.450849

**Authors:** Shiraz A. Shah, Ling Deng, Jonathan Thorsen, Anders G. Pedersen, Moïra B. Dion, Josué L. Castro-Mejía, Ronalds Silins, Fie O. Romme, Romain Sausset, Eric Olo Ndela, Mathis Hjemlsø, Morten A. Rasmussen, Tamsin A. Redgwell, Gisle Vestergaard, Yichang Zhang, Søren J. Sørensen, Hans Bisgaard, Francois Enault, Jakob Stokholm, Sylvain Moineau, Marie-Agnès Petit, Dennis S. Nielsen

**Author notes:** These authors contributed equally (Shiraz A. Shah, Ling Deng).

## Abstract

The gut microbiome (GM) is shaped through infancy and plays a major role in determining susceptibility to chronic inflammatory diseases later in life. Bacteriophages (phages) are known to modulate bacterial populations in numerous ecosystems, including the gut. However, virome data is difficult to analyse because it mostly consists of unknown viruses, i.e. viral dark matter. Here, we manually resolved the viral dark matter in the largest human virome study published to date. Fecal viromes from a cohort of 647 infants at 1 year of age were deeply sequenced and analysed through successive rounds of clustering and curation. We uncovered more than ten thousand viral species distributed over 248 viral families falling within 17 viral order-level clades. Most of the defined viral families and orders were novel and belonged to the *Caudoviricetes* viral class. Bacterial hosts were predicted for 79% of the viral species using CRISPR spacers, including those in metagenomes from the same fecal samples. While *Bacteroides*-infecting Crassphages were present, novel viral families were more predominant, including phages infecting Clostridiales and *Bifidobacterium*. Phage lifestyles were determined for more than three thousand caudoviral species. Lifestyles were homogeneous at the family level for 149 *Caudoviricetes* families, including 32 families that were found to be virulent, while 117 were temperate. Virulent phage families were more abundant but temperate ones were more diverse and widespread. Together, the viral families found in this study represent a major expansion of existing bacteriophage taxonomy.

## Introduction

The establishment of the gut microbiome (GM) during the first years of life plays a pivotal role in the maturation of the infant immune system^1, 2^. Early-life GM dysbiosis has been linked to a series of chronic diseases occurring later in life, indicative of an immune system thrown off balance^3–6^. Most existing research has been on the bacterial component of the GM but in recent years it has become evident that other microbes as well as viruses are prominent GM members. The latter colonize the gut during the first months of life following a patterned trajectory resembling the establishment of gut bacteria^7–10^.

Bacteriophages (phages) are viruses that infect bacteria in a host specific manner. Virulent phages multiply by killing their host. Temperate phages can integrate their genome into the bacterial chromosome, thereby becoming prophages and postponing an attack on the host until conditions are favourable. Some phages also cause chronic infections leading to continuous shedding of viral particles^11^. Bacteria will defend themselves using an impressive arsenal of defence systems, which include among others, CRISPR-Cas systems, an adaptive immune mechanism where DNA records (spacers) of past infections are saved on the bacterial CRISPR array to help combat any future phage attacks^12^.

Lately, it has become clear that phages possess the ability to alter GM composition and function^8, 13^. Moreover, the reported interactions between phage proteins and the host immune response^14–16^ suggest a tripartite interaction that may modulate host health. The first report on the viral metagenome (virome) composition in the infant gut dates back more than a decade^17^, and it has recently been shown to be influenced by caesarian section^18^. Nevertheless large-scale studies establishing the early-life virome composition and structure are sparse, and human virome studies in general have been challenged by the viral “dark matter” problem^19^.

The dark matter problem is a phenomenon where only a small fraction of nucleic acid sequences can be traced back to any known virus because sequence databases currently under-represent the diversity of viruses in natural environments. Attempts at *de novo* virus discovery directly from the virome data have been limited by the lack of universal viral marker genes, which makes it difficult to distinguish viral sequences from contaminating DNA. In addition, *de novo* classification of novel viruses has been held back by the lack of criteria and standardised methods. Although much progress has been made in recent years^20–22^, virome studies are still catching up. Thus, the dark matter problem has prevented researchers from getting the most out of their viromes, perhaps thereby missing biologically meaningful associations in their data. We posit that comprehensive annotation of viromics data, including viral taxonomies, trees and gene families will be required for more powerful statistics against sample metadata. Such an effort would inevitably involve cataloguing novel viruses, and ideally proposing novel viral taxa so they can be known to science.

Traditionally, the definition of new viral taxa has required the laboratory isolation of the virus along with its host and its subsequent characterization^23^. Recently the International Committee for the Taxonomy of Viruses (ICTV) opened up the possibility for defining viral taxa based on sequence information alone. This move is due to have major implications for our understanding of phages, and tailed phages (or caudoviruses) in particular, as their diversity is the greatest and has been misrepresented so far^24^. As an example, the ICTV established the complete taxonomy of the new *Herelleviridae* caudoviral family, demonstrating the proper definition of viral subfamilies and genera under the new paradigm^25^. Subsequently, three new caudoviral families were found in human gut metagenome data^26^, and recently the prominent human gut phage family *Crassviridae*^27^, was elevated into a viral order *Crassvirales*^28^, belonging to the viral class *Caudoviricetes* which itself is proposed to encompass caudoviruses^29^ in general.

Here, we present the characterization of the fecal virome from 647 infants at one year of age enrolled in the COPSAC2010 cohort^30^. The viral dark matter was exhaustively resolved by manual curation, which enabled the identification of over ten thousand viral species, more than half of which appear to be completely sequenced. Hierarchical clustering of these viruses based on encoded protein similarity enabled the *de novo* definition of novel viral genera, subfamilies and families. In total, we identified 248 viral families that fall within seventeen viral order-level clades. Most of the novel families belonged to *Caudoviricetes*, representing a major expansion of known caudoviral diversity. We also predicted the hosts of the viral species by matching spacer sequences from bacterial CRISPRs and found that members of the *Bacteroides*-infecting *Crassvirales* - otherwise abundant in adult gut viromes - were outnumbered by novel phage families infecting numerous gut bacteria such as Clostridiales and *Bifidobacterium*. Temperate phages were the most widespread and diverse while virulent phages were more abundant. Our manually confirmed viral set was also used for benchmarking the performance of several metagenome virus discovery tools. All viral sequences found in the study have been made available online in an interface for browsing their genome contents along with their viral taxonomy, host and lifestyle predictions (http://copsac.com/earlyvir/f1y/fig1.svg).

## Results

### Study population

COPSAC2010 is a population-based birth cohort of 700 children recruited in pregnancy where we study the causes for chronic inflammatory diseases. The children are monitored by clinicians who diagnose eventual conditions, and multi-omics data is collected routinely to shed light on disease mechanisms. A total of 647 children had a fecal sample obtained at 1 year of age where the virome was characterised, and deep metagenomes for the samples were sequenced in parallel^31^.

### Identifying the viruses and resolving their taxonomies

Virome extractions are known to contain large amounts of bacterial contaminating DNA^32^ and the virome dark matter problem^19^ makes it difficult to discern novel viruses from contaminants. We elected to resolve the virome dark matter by assembly, clustering and successive rounds of manual curation.

Assembly of the 647 virome samples resulted in 1.5M contigs larger than 1kb that fell within 8050 broad decontamination clusters based on protein similarity (Figure S1). After ranking the clusters by prevalence and CRISPR targeting, they were visualised as in Figure S2 and inspected manually. A total of 255 clusters (3.2%) were deemed viral. In parallel, we deduplicated all 1.5M contigs at the species level, (95% average nucleotide identity or ANI), resulting in 363k operational taxonomic units (OTUs) (Figure 2). Of these, 16,746 belonged to the 255 viral decontamination clusters and were thus termed viral OTUs (vOTUs).

The vOTUs were pooled with all 7.7k available reference phage species^93^ before gene calling. Protein alignments were then used for defining viral ortholog gene clusters (VOGs) *de novo* and for constructing an aggregate protein similarity (APS) tree. The tree was rooted and cut at the levels reproducing the recent taxonomy for the *Herelleviridae*^25^ phage family, thus yielding viral families, subfamilies and genera covering all vOTUs and reference phages. An additional order-level cutoff was based on the newly proposed caudoviral *Crassvirales* order^28^.

All family-level clades were validated by visualising (Figure S2) and inspecting their gene contents and any weak clades were removed in the process. The minimum complete genome size cutoff was determined by examining the vOTU size distribution within each family. Next, each vOTU within each family was curated individually to remove 6,725 vOTUs comprising small fragments, putative satellites and MGEs.

The final curated set consisted of 248 viral families, including sixteen known families and 232 novel ones (Figure 1). The novel viral families were named after the infants that delivered the fecal samples. All the families were additionally grouped into seventeen order-level clades, twelve of which were novel (Table 2). The 248 curated families harboured 10,021 manually confirmed species-level vOTUs, 5,608 of which were complete or near-complete. DNA sequences and taxonomies for the vOTUs along with visualisations of the families (Figure S2) have been made available online via the interactive version of Figure 1 at http://copsac.com/earlyvir/f1y/fig1.svg

**Figure 1:**
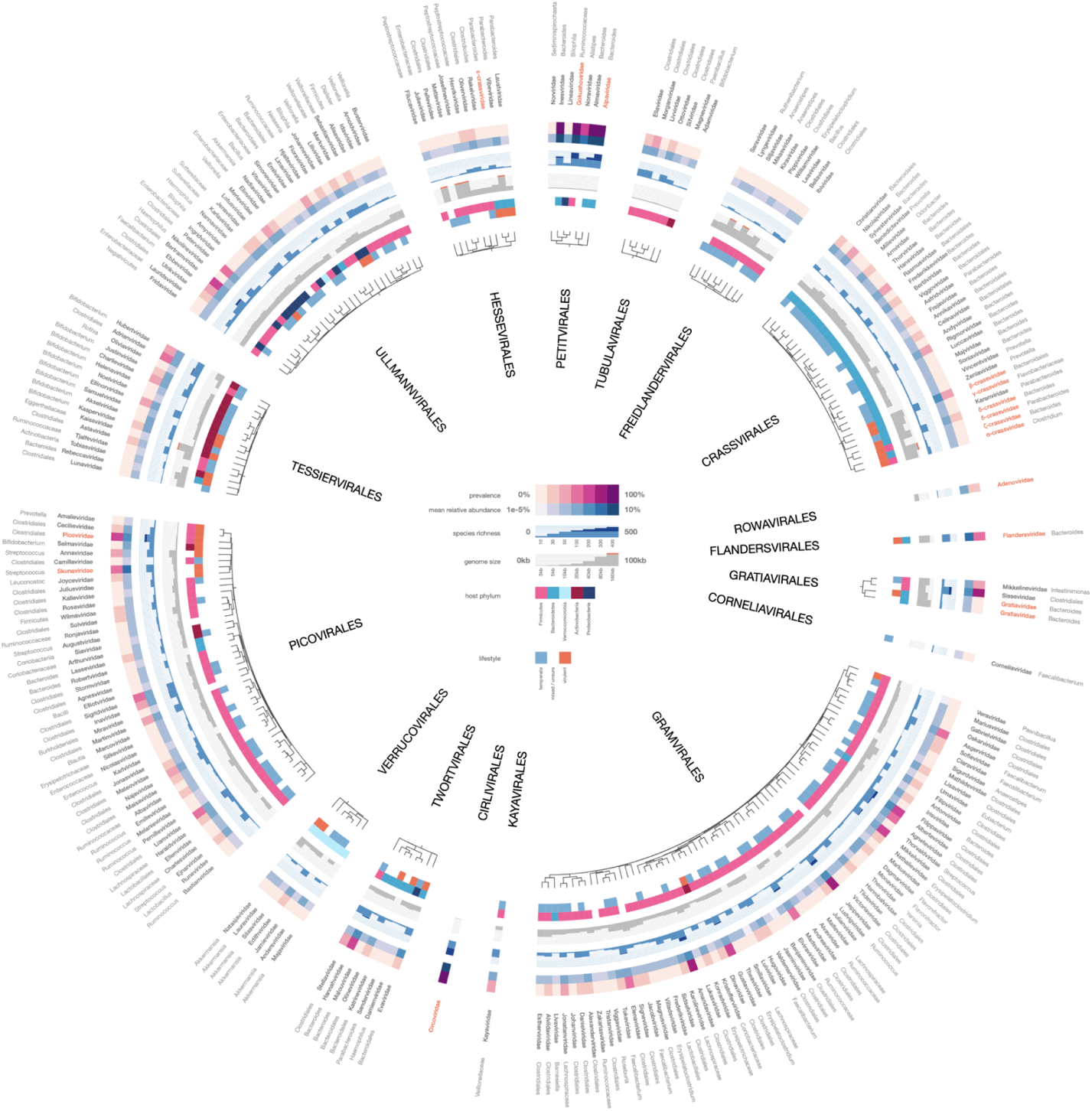
An atlas of infant gut DNA virus diversity. Fecal viromes from 647 infants at age 1 year were deeply sequenced, assembled and curated, resulting in the identification of 10,021 viruses falling within 248 viral families. Predicted host ranges for each family are given, and the families have been grouped into 17 order-level clades. Trees show how families are interrelated within each order-level clade. The 16 previously defined families are highlighted in red. An interactive version with expandable families can be accessed online: http://copsac.com/earlyvir/f1y/fig1.svg

### vOTU host distributions mimic sample bacterial composition

Bacterial hosts for the vOTUs were predicted using 318k CRISPR spacers from our metagenome assembled genomes (MAGs)^31^ and the 11M spacers from the CRISPR spacer database^33^ as well as by using WIsH^34^. The three host predictions were merged by selecting the last common ancestor (LCA). 63% of the vOTUs yielded host predictions at the bacterial genus-level, while 77% were predicted at the bacterial order-level (Figure 3A) and 79% at the phylum level. *Bacteroides* was by far the most common bacterial host genus followed by *Faecalibacterium* and *Bifidobacterium*. At the order level, more than half of all vOTUs had Clostridiales as hosts, with Bacteroidales covering just 20% (Figure 3A). These differences mirror the corresponding pattern for the bacteria found in the metagenome, where *Bacteroides* are abundant while Clostridiales are diverse (Figure 3B).

### Quantification of viruses and bacterial contamination

In order to gauge the quality of the virome extractions we estimated the virus particle concentration for a subset of the samples using epifluorescence microscopy. The mean virus-like particle (VLP) concentration obtained was 1.0 × 10^9^ VLPs/g of feces, ranging from 3 × 10^8^ to 3 × 10^9^ VLPs/g for the 18 samples tested.

Before library preparation we used multiple-displacement amplification (MDA) because it enables the detection of ssDNA viruses. MDA, however, can introduce compositional biases^35, 36^ that favour ssDNA viruses over dsDNA viruses. To limit any bias we kept the MDA step at 30 minutes instead of the recommended 2 hours. After sequencing and assembly, vOTU abundances were estimated by read mapping and normalising for mapping depth and contig length. ssDNA viral counts made up more than half of the mean relative abundance (MRA) across all samples. To investigate if the MDA had biased the caudoviral counts, we compared counts of plaque forming units (PFUs) of different coliphages isolated from the same samples^37^ against their corresponding vOTU abundances. Phages that were present in titers lower than 25000 PFU/g of feces were undetected, limited most likely by sequencing depth. However, phages with PFU counts above this limit showed reliable quantitative abundances (Figure S3). This confirmed that the virome abundances were quantitative for caudoviruses internally, allowing for valid comparisons of dsDNA abundances across different vOTUs and samples.

The 10,021 confirmed vOTUs recruited roughly half of the reads despite making up a minor fraction of all OTUs (Figure 2). The remaining half of the read mappings were spread over the 346k non-vOTUs. Since ViromeQC^38^ estimated a mean bacterial contamination rate of 44%, we infer that most non-vOTUs must be bacterial (Figure 2). After subtracting viral and bacterial reads, only 7% of the reads were left over as unaccounted dark matter.

**Figure 2:**
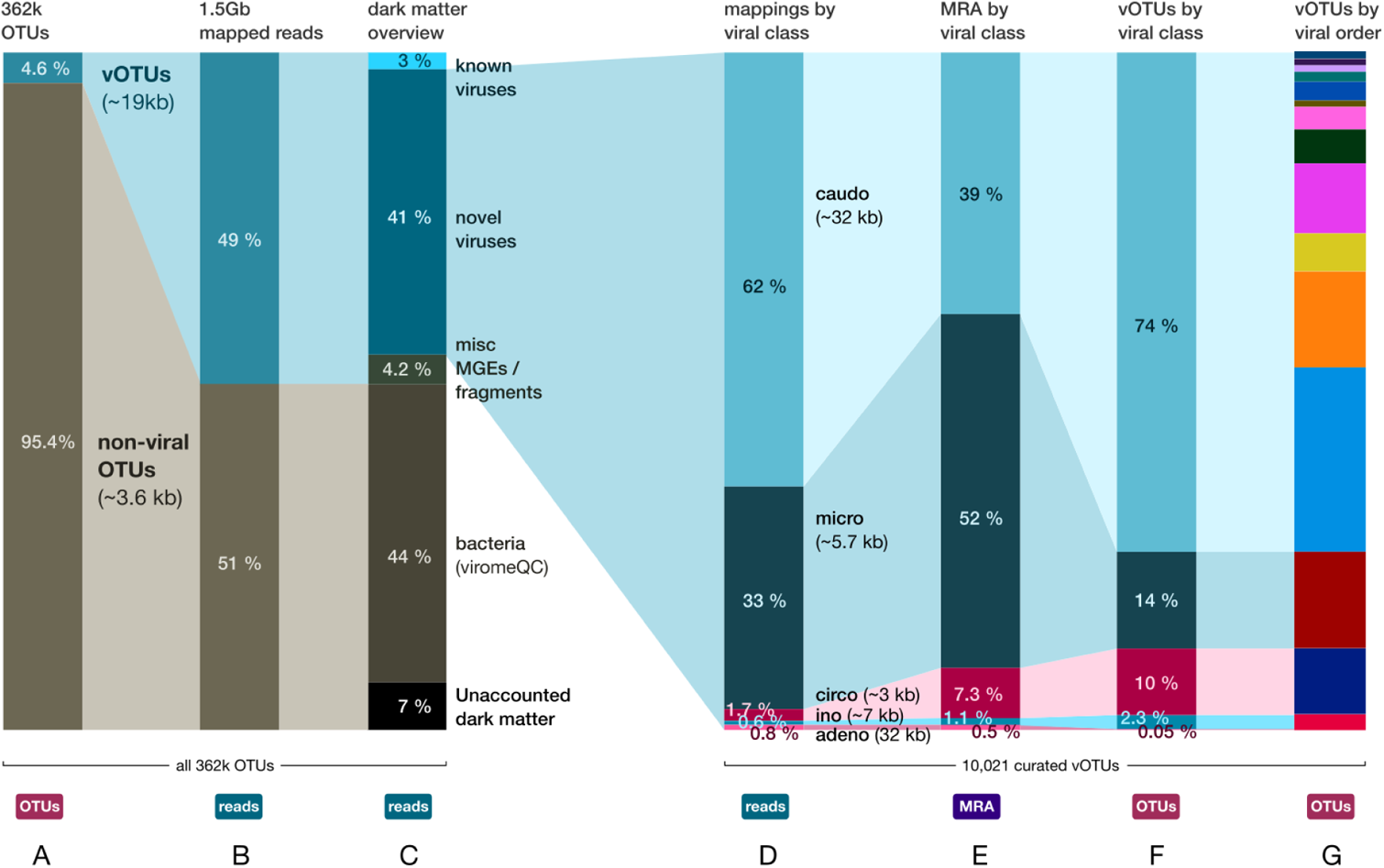
From assembly to curated vOTUs in numbers. After assembly, species-level deduplication and manual decontamination, most OTUs were found to be non-viral and had small sizes while viral OTUs were much fewer but longer (A). After mapping, vOTUs accounted for roughly half of the reads (B). 97% of the reads originally comprised dark matter but only 7% was left after resolution (C). The 10,021 curated vOTUs fell within five viral classes (caudoviruses, microviruses, circoviruses, inoviruses and adenoviruses). Distributions of the viral classes by: mapped reads (D), mean relative abundances (E) and species richness, i.e. number of vOTUs (F) are shown. G) Same as F but at viral order-level, with orders colored as in Figure 4.

**Figure 3:**
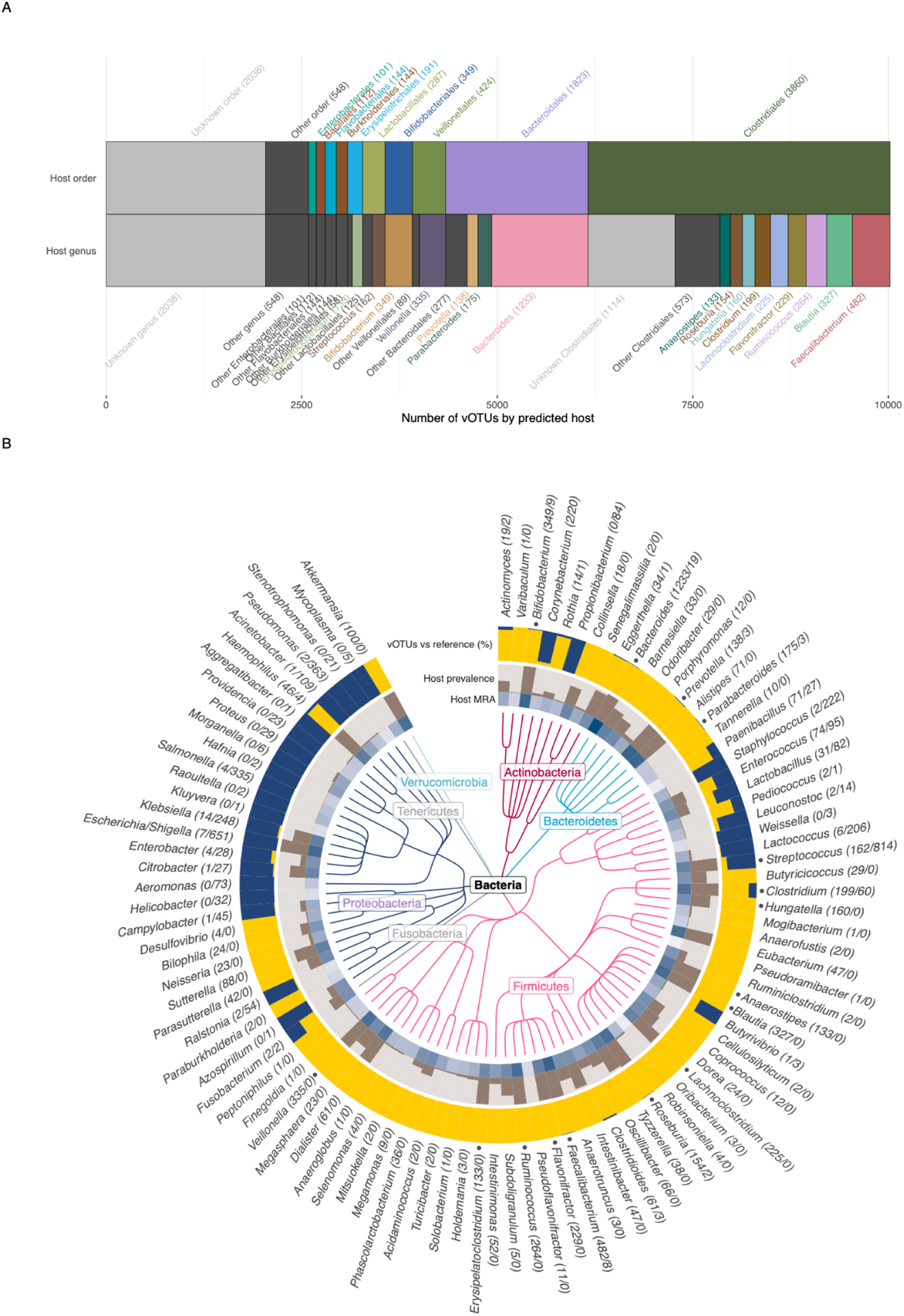
Prediction of bacterial hosts for the 10,021 viruses found in the infant gut virome shows that *Bacteroides*, *Faecalibacterium* and *Bifidobacterium* are the three most prominent host genera. A) Distribution of virus host predictions collapsed to bacterial order and genus levels, respectively. Numbers in parentheses denote the number of vOTUs with a given host genus or order. B) The top 100 gut bacterial genera found in gut metagenomes from the same infant fecal samples, as represented by a taxonomic tree. The mean relative abundance (MRA) of each bacterial genus is shown in the blue heatmap, while the fraction of the 647 infants harbouring the host genus (i.e. its prevalence) is shown with the brown barplot. For each host bacterium, the yellow bars show the proportion of viral species found in this study relative to those in the reference phage species set^91^ shown in dark blue. Numbers behind each genus name denote the total number of vOTU vs. reference phage species per bacterial host genus. The 16 major host genera from panel A are indicated by a dot in front of their names in panel B.

### Viral family prevalence, richness and abundance

In order to identify the predominant viral families, we ranked them by both species richness, prevalence across samples, and MRA (Figure 4). All three rankings gave similar results because the three estimates were highly correlated (Figure 1). The correlation between these measures is predicted by neutral theory which has already been shown to explain bacterial community structures quite accurately^39, 40^. Human-infecting ssDNA circoviruses (*Cirlivirales*) and bacterial ssDNA microviruses (*Petitvirales*) were the most abundant (Figure 4A). The abundant ssDNA viral families were followed by the top ten most abundant major caudoviral families (Figure 4B). Of the major caudoviral families, four were already known, namely *Skunaviridae*, *Flandersviridae*, *Picoviridae* and *β-crassviridae*, while the remaining six were novel.

**Figure 4:**
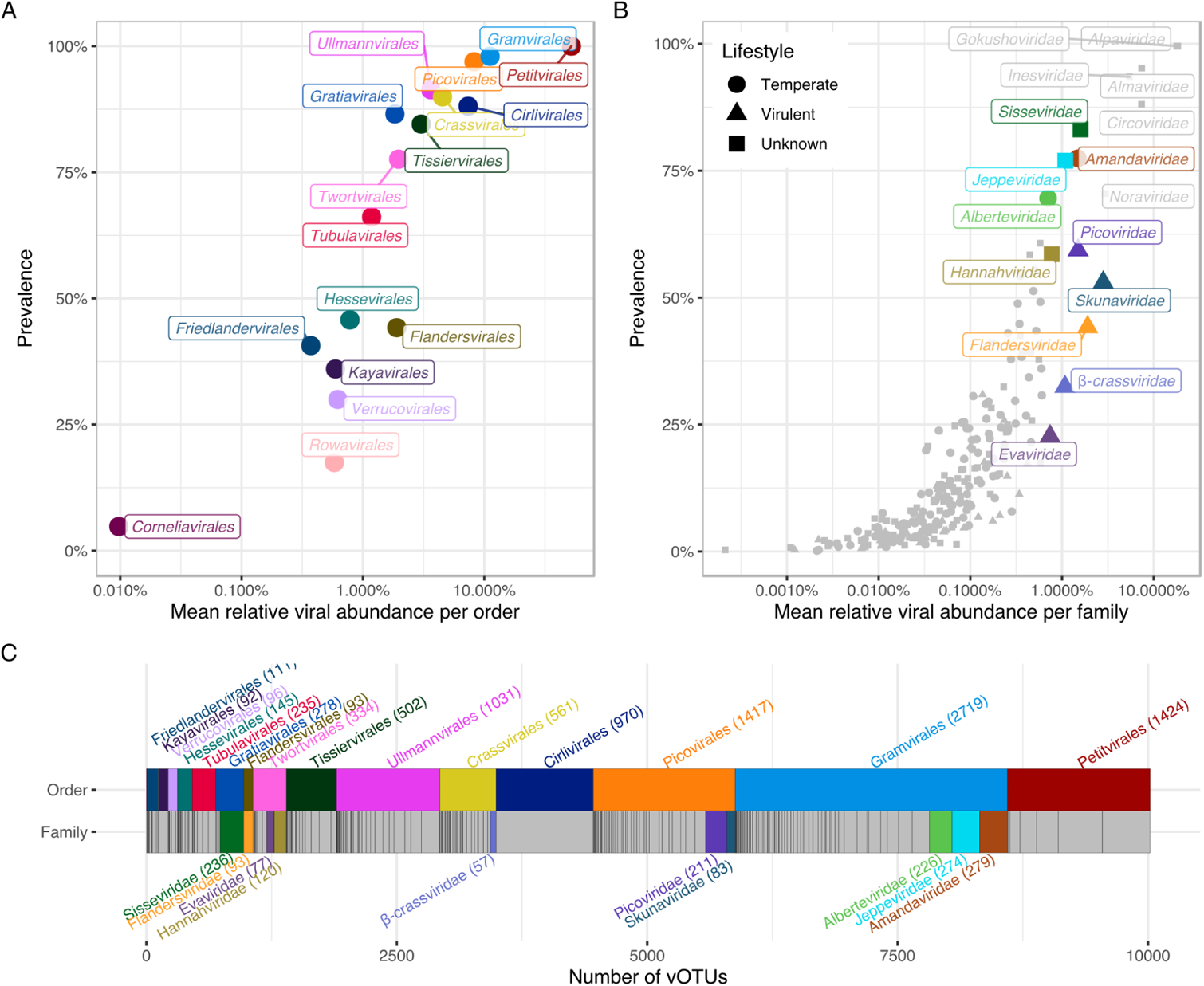
Abundance, prevalence and richness of the novel families in the infant gut. A) The 17 viral order-level clusters and their prevalence and mean relative abundance (MRA) across samples. B) The 248 viral families in terms of prevalence and MRA. The major caudoviral families are colored and labelled. Minor families as well as ssDNA families are in grey. Predicted lifestyles for the 10 major caudoviral families are indicated by different shapes. C) Viral orders and families scaled by species richness, ordered by MRA. The viral families are represented underneath the order they belong to. The major families were defined as the ten most abundant caudoviral families in the data.

### Virulent vs. temperate *Caudoviricetes* families

While examining the ten most abundant major caudoviral families we noticed that most of them lacked an encoded integrase, otherwise commonly found throughout the less abundant families in the data. Since an integrase is an indicator of a temperate lifestyle, we went on to investigate systematically whether a virulent lifestyle was linked to higher abundances overall. Using the determined minimum complete size limit per viral family, 3398 complete and near-complete caudoviral vOTUs from 230 caudoviral families were screened for the presence of integrase VOGs. Using this information 117 of the caudoviral families were deemed temperate, while only 32 were found to be virulent. The remaining 81 caudoviral families exhibited either a mixed lifestyle pattern or were uncertain due to an insufficient number of complete genomes.

On its own, abundance was not significantly linked to phage lifestyle (Figure 5A), but temperate families were significantly more prevalent than virulent families (Figure 5B). Temperate phages have been shown to be more diverse than their virulent counterparts^41^, so we looked into this by comparing their genus richness normalised for overall family size. Indeed temperate caudoviral families were significantly more genetically diverse than virulent families (Figure 5C). As for the predicted bacterial hosts, *Clostridiales* were particularly enriched in temperate viral families, whereas most virulent families were predicted to infect *Bacteroidales* (Figure 1). Using the CRISPR spacer mappings we found, in line with observations in other studies^22, 42^, that some vOTUs appeared to infect multiple host species, genera or even families of bacteria. We decided to check whether the CRISPR-Cas system targeted virulent phages more often than temperate phages, or whether virulence was associated with a broader host range. This was not the case as both temperate and virulent families exhibited similar mean host ranges and numbers of targeting spacers (Figure 5DE).

**Figure 5:**
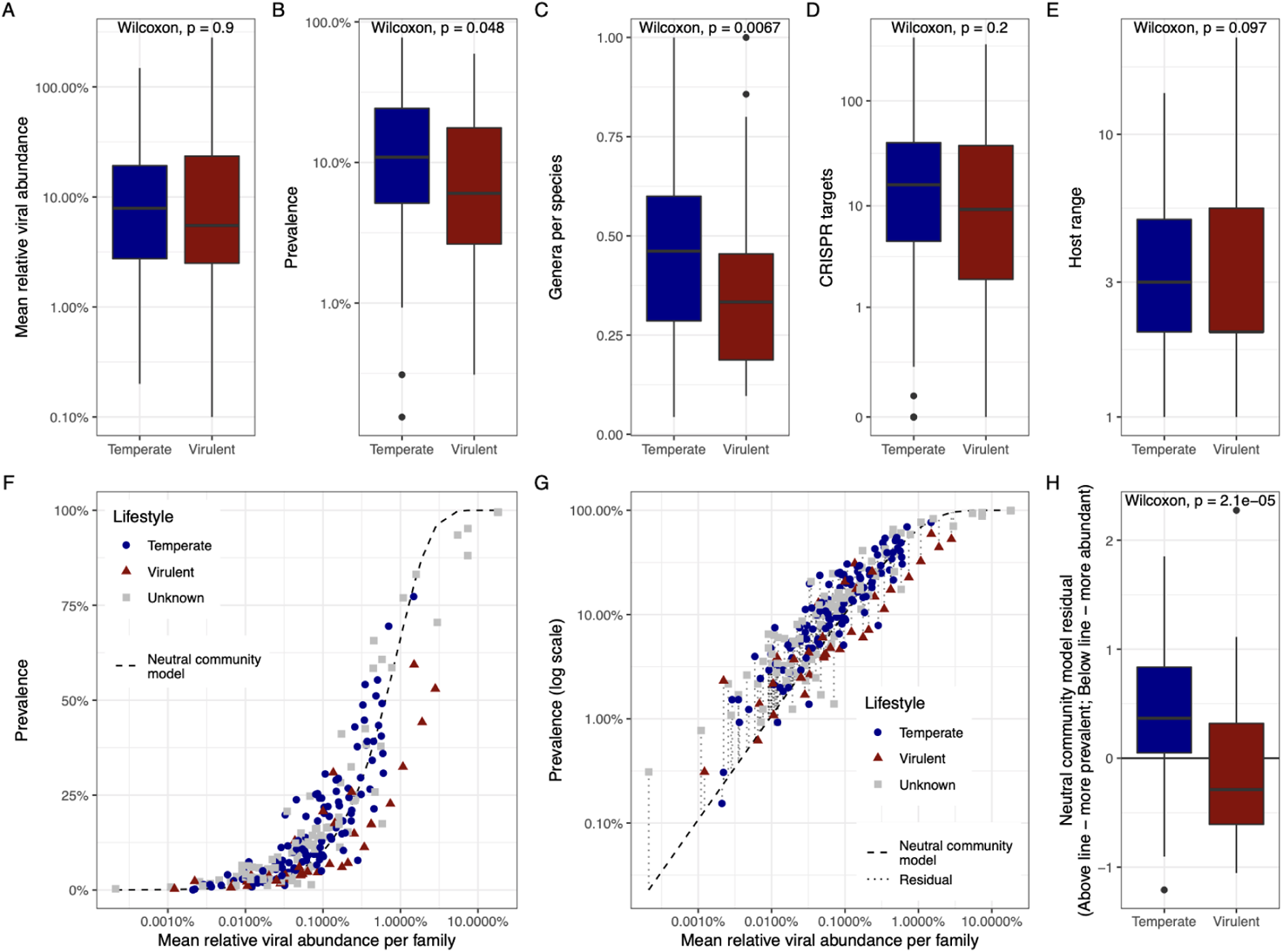
Characteristics of temperate vs. virulent families in the data in terms of A) Mean relative abundance; B) Prevalence; C) Genus richness for family size; D) Number of metagenomic CRISPR spacer matches; E) Host range (number of host species); F) Fit of the neutral community model on the viral families from Figure 4B, with families coloured by lifestyle, G) Deriving neutral community model residuals from the log-transformed prevalences; and H) Comparison of neutral community model residuals, showing that temperate families tend to have positive residuals, whereas virulent families tend towards negative residuals.

Plotting the abundance and prevalence of the virulent and temperate families against each other (Figure 5F) resurfaced our initial suspicion that virulent phage families were present in higher numbers despite being found in fewer children. Thus we decided to test this hypothesis systematically by using the neutral community model (Figure 5G), which describes the typical relationship seen in nature between abundance and prevalence, as the baseline assumption^43^. After fitting the model on all of our family abundances, temperate families had significantly higher residuals against it than virulent families (Figure 5H), confirming that they were more prevalent while also being less abundant than virulent phage families.

### ssDNA viruses in the infant gut

ssDNA vOTUs collectively recruited around a third of the sequencing reads, but after normalising for their short genome sizes, they accounted for 60% of the MRA (Figure 2). Although MDA may have inflated the counts for ssDNA viruses, both their prevalence and richness was in line with their high abundance. The ssDNA virus families display canonical positioning along the neutral community model (figures 4B, 5F) indicating that any inflation should have been limited. ssDNA viruses in our data fell within three separate viral classes, *Malgrandeviricetes*, *Arfiviricites* and *Faserviricetes*, harboring a single viral order each.

Microviruses of the *Petitvirales* viral order (class *Malgrandeviricetes*) were both the most prevalent and abundant group of viruses found in our viromes, making up 52% of the MRA. Further, 21 % of the CRISPR spacer matches from the metagenome targeted microviruses. This was in line with their overall richness which accounted for 16% of all bacterial viruses in our data, or 1424 vOTUs in total. vOTUs from the two major families, *Gokushoviridae* and *Alpaviridae* (currently known as *Gokushovirinae* and *Alpavirinae*) in our data infect Clostridiales and Bacteroidales respectively, but other minor and novel microviral families were also detected.

*Circoviridae*, also referred to as anelloviruses, is a single family of small 3-kb ssDNA viruses that infect animal tissue. They are known to cause chronic asymptomatic infections in humans, displaying elevated titers in individuals with weak or developing immune systems^44^. The immature immunity of the infants may explain why *Circoviridae* were so abundant in our samples, making up 7% of the MRA and comprising by far the richest single family with 970 species-level vOTUs. On average, each infant harboured 10 different species of *Circoviridae*. Unsurprisingly, no CRISPR spacer matches were found targeting any *Circoviridae* vOTUs.

Inoviruses from the *Tubulavirales* order are small ssDNA filamentous phages recently found to be ubiquitous and diverse^45^. Some of them can integrate into their host genomes using encoded integrases while others cause chronic non-lethal infections that are neither lytic, nor lysogenic, but result in the continuous shedding of new virus particles^11^. Although they were diverse in our data, split into 7 distinct families, like the *Petitvirales*, their species richness was much lower at 235 vOTUs, and abundances were correspondingly lower at 1% MRA. Most of the inoviral families found, were predicted to infect Clostridiales, although members of the *Adamviridae*, appear to specifically infect *Bifidobacterium*.

### The major *Caudoviricetes* families

As explained previously, viral families were defined here by cutting the APS tree at the branch uniting *Herelleviridae*^25^, and this cutoff was validated independently because it reproduced the expected Crassphage families just recently defined^28^. Ranking the 230 obtained caudoviral families by MRA, the virulent *Skunaviridae* was the most abundant caudoviral family in the children, comprising 2.7% MRA overall or 6.3% when counting only caudoviruses and disregarding ssDNA viruses. Our most abundant *Skunaviridae* vOTUs resemble numerous reference phage genomes infecting *Lactococcus* dairy cultures. However, most vOTUs from this family were predicted to infect *Streptococcus*, which was a highly prevalent bacterial host in our samples. *Flandersviridae*, a recently described family of gut phages^26^ that infects *Bacteroides*, was the second most abundant caudoviral family in the children at just under 5.9% caudoviral MRA. Consistent with earlier speculation^26^, we predicted this family to be virulent. Next, at 3.9% caudoviral MRA was our first novel viral family, which we named *Sisseviridae*. It was also the most prevalent caudoviral family, found in more than 80% of the samples. This family included the recently discovered *Faecalibacterium* phage Oengus^46^, also known to be highly prevalent. Being a large family composed of 236 vOTUs infecting mainly Clostridiales hosts, it displayed a mixed lifestyle with some subfamilies being virulent and others temperate. The fourth most abundant caudoviral family was the *Picoviridae* at 3.8% of the caudoviral MRA. This family comprises a virulent group of podoviruses including reference phages such as *Bacillus* phage phi29. The vOTUs belonging to this family were split between those infecting Actinobacteria such as *Bifidobacterium* and *Eggerthella*, or Clostridiales hosts like *Erysipelatoclostridium* and *Hungatella*. At 3.5% caudoviral MRA, was our most abundant Crassphage family, *β-crassviridae*. Of note, this family is different from the *α-crassviridae* most commonly found in adults^47^. The latter was only present at 0.2% caudoviral MRA in our infant samples. The predicted hosts for *β-crassviridae* included both *Clostridium* and *Bacteroides*. Next, *Amandaviridae* and *Jeppeviridae* at 3.2 and 2.7% caudoviral MRA comprise two related and large families of temperate phages containing almost three hundred vOTUs each, infecting Clostridiales host genera such as *Ruminococcus*, *Blautia*, *Anaerostipes* and *Hungatella*. Both families share the interesting characteristic of terminase shuffling, where the conserved caudoviral gene, *terL*, is subject to frequent horizontal gene transfer (HGT), while the remainder of the genome is held relatively constant. The related family *Alberteviridae* exhibits a similar characteristic. Finally, *Evaviridae* and *Hannahviridae* at 2.4 and 2.3% caudoviral MRA, respectively, comprise two related novel families of *Bacteroides*-infecting phages. The former was predicted to be virulent while the latter was mixed containing two major subfamilies each with its own lifestyle. *Hannahviridae* includes the recently described *Bacteroides* Hankyphage^48^, and has been extensively described in our parallel provirome study performed on the same samples^94^.

### Novel orders within *Caudoviricetes*

Cutting the APS tree at the branch unifying the recently defined *Crassvirales* viral order reproduced all other known viral orders in the virome, including *Petitvirales* (microviruses), *Tubulavirales* (inoviruses), *Cirilvirales* (circoviruses) and *Rowavirales* (adenoviruses). In addition to the known orders, the cutoff predicted the existence of at least 12 novel viral orders all of which were caudoviral (Table 1), unifying our families into broader groups of related families. *Gramvirales*, the most diverse, prevalent and abundant clade at 27% caudoviral MRA, encompassed 67 novel viral families that were overwhelmingly temperate, infecting Clostridiales host bacteria. *Gramvirales* includes the major families *Jeppeviridae*, *Amandaviridae* and *Alberteviridae* described above. Interestingly, terminase shuffling was a feature that seemed to pervade *Gramvirales*. Two major TerL clades are prevalent throughout most of the order, and they are often exchanged even between species of the same genus. Some caudoviral families were so distantly related to any other family that they formed their own singleton orders. *Flandersvirales* is a notable example but we found others as well, like *Kayavirales*, composed of a single novel family *Kayaviridae* that infects mostly *Veillonella*. Our *Crassvirales* clade includes the four original crassphage families^49^ plus the recently described *ζ-crassviridae*, which were all predicted to be virulent. In addition, 24 mostly temperate families, infecting mainly *Bacteroides*, were also found to be part of the same order. However, *ε-crassviridae* belongs to a different order-level clade, *Hessevirales*, consisting of multiple families with large genomes. The *Crassvirales* clade as a whole covers 15% of the caudoviral MRA, with the previously proposed families making up 6%. Other caudoviral families overshadow Crassphage families in our data, and *Crassvirales* as an order is less prevalent than the novel *Gramvirales*, *Picovirales* and *Ullmannvirales*.

**Table 1:** The 10,021 viral species belonging to the 248 viral families found in the present study were grouped into viral order-level clades. The clades are sorted by total richness and novel orders are indicated in bold. Caudoviral MRAs (cMRA) are shown for the caudoviral orders. For each order-level clade, the number of families, subfamilies and genera are also given along with their most frequent hosts.

| order-level cluster | % cMRA | families | subfams | genera | species | major host |
| --- | --- | --- | --- | --- | --- | --- |
| **Gramvirales** | 27 | 67 | 531 | 1163 | 2719 | Clostridiales |
| *Petitvirales* | - | 7 | 67 | 154 | 1424 | *Bacteroides* |
| **Picovirales** | 19 | 45 | 315 | 618 | 1417 | Clostridiales |
| **Ullmannvirales** | 9 | 30 | 171 | 358 | 1031 | Veillonellaceae |
| *Cirlivirales* | - | 1 | 21 | 268 | 970 | Human |
| *Crassvirales* | 15 | 30 | 114 | 214 | 561 | Bacteroidales |
| **Tissiervirales** | 7 | 17 | 64 | 149 | 502 | *Bifidobacterium* |
| **Twortvirales** | 6 | 8 | 47 | 108 | 334 | Bacteroidales |
| **Gratiavirales** | 5 | 4 | 27 | 75 | 278 | Clostridiales |
| *Tubulavirales* | - | 7 | 50 | 78 | 235 | Clostridiales |
| **Hessevirales** | 2 | 11 | 37 | 61 | 145 | Clostridiales |
| **Friedlandervirales** | 1 | 10 | 38 | 61 | 111 | Clostridiales |
| **Verrucovirales** | 1 | 7 | 19 | 31 | 96 | *Akkermansia* |
| **Flandersvirales** | 6 | 1 | 4 | 10 | 93 | *Bacteroides* |
| **Kayavirales** | 1 | 1 | 3 | 4 | 92 | Veillonellaceae |
| **Corneliavirales** | 0.05 | 1 | 3 | 6 | 8 | *Faecalibacterium* |
| *Rowavirales* | - | 1 | 1 | 2 | 5 | Human |

### Benchmarking virome decontamination software

A series of virus discovery and decontamination tools have been published recently^21, 38, 50–56^ and they have already seen widespread application by the viromics community. Several large gut phage databases have been released that were built on predictions from such tools^22, 36^. Yet, little is known about their efficacy in identifying novel viral clades or their ability to weed out contaminating DNA. The manually curated nature of our virome data set made it well suited for independently testing the performance of these tools. A purely random prediction was generated for comparison. A naive length cutoff of +20kb was also used for comparison, since non-vOTUs were distinctly short in our data (Figure 2).

CheckV^50^, VIBRANT^21^ and viralVerify^52^ sported the best performances with our data set (Table 2) although VIRSorter^55^ also worked well. With a specificity of 97.5%, the length cutoff did a better job than VIRSorter2^51^. DeepVirFinder^56^ PPR-Meta^54^ and Seeker^53^, all of which were “alignment free”, yielded performances that were close to random. VIRSorter and VIBRANT, when run in virome decontamination mode, improved sensitivity at the cost of greatly reduced specificity. For our data, VIRSorter performed better when used in “db2” mode.

**Table 2:** Benchmarking statistics from various metagenomic virus discovery methods against 10,021 vOTUs in the manually curated viral set from a total of 362,668 OTUs. The skewed nature of the data set, with non-vOTUs far outnumbering vOTUs, inflates the specificity metric, which is 0.965 for a random prediction. The Kappa performance metric was included because it is robust against skews. Prediction methods that provided a confidence score were cut conservatively or by matching the number of positive predictions to the manual set. The performances of VIRSorter and VIBRANT were also checked in alternate modes (bottom portion).

| method | predictions | cutoff | sensitivity | specificity | kappa |
| --- | --- | --- | --- | --- | --- |
| CheckV | 9393 | >= medium qual | 0.71161 | 0.99359 | 0.7273 |
| viralVerify | 10921 | > 15 | 0.69873 | 0.98889 | 0.6589 |
| VIBRANT | 5843 | >= medium qual | 0.44696 | 0.99613 | 0.5556 |
| VIRSorter | 14366 | cat. 1 + 2 | 0.69335 | 0.97896 | 0.5553 |
| Length + 20kb | 13452 | > 20kb | 0.48548 | 0.97565 | 0.3954 |
| VIRSorter2 | 18638 | virus maxscore = 1 | 0.50983 | 0.96164 | 0.3325 |
| DeepVirFinder | 10192 | q < 0.05 | 0.058078 | 0.972749 | 0.0306 |
| PPR Meta | 17222 | score > 0.95 | 0.060772 | 0.952891 | 0.0101 |
| random | 12500 | w/o replacement | 0.0330306 | 0.9654924 | -0.0013 |
| Seeker | 13520 | >= 0.9 | 0.0263447 | 0.9624100 | -0.0096 |
| VIRSorter virome | 30736 | cat. 1 + 2 | 0.83155 | 0.93647 | 0.3832 |
| VIRSorter db2 | 13298 | cat. 1 + 2 | 0.70971 | 0.98246 | 0.5973 |
| VIBRANT virome | 6181 | >= medium qual | 0.45395 | 0.99537 | 0.5521 |

ViromeQC estimates the proportion of bacterial contamination in viromes^38^ and it estimated that around half of our reads were contaminants. We compared the ViromeQC estimation to two other independent measures that we generated using the manual curation and our coupled metagenome samples, namely 1) the proportion of mapped reads to non-vOTUs and 2) the depletion of bacterial core genes in the virome reads compared to cognate metagenomes. Although all three estimates were in strong agreement on average, there was considerable sample to sample variation (Figure S4). Importantly, our metagenome-normalised core gene depletion did not perform better than ViromeQC when comparing against the non-vOTU standard. This result illustrates that virome contaminant estimation is non-trivial and that ViromeQC performs well.

## Discussion

We have presented here the largest human virome study published to date, covering fecal samples from 647 1-year-old infants where DNA viromes were sequenced to an average depth of 3 Gb per sample. This study additionally represents three major advances in descriptive viromics. To our knowledge, this has been the first exhaustive attempt at resolving virome dark matter. We were able to match 93% of the reads to either viral or bacterial DNA, leaving only 7% as unaccounted dark matter. Secondly, we performed the first reported comprehensive taxonomic resolution of a virome data set. This led to the identification of 232 novel viral families, representing a major expansion of known phage taxonomy. Lastly, we generated an overview of phage lifestyles for an entire ecosystem. The analysis shows that the bulk of phage diversity in the infant gut is composed of temperate phages, even if less diverse virulent phages can be more abundant.

Bacterial contaminating DNA made up just under half of our sequenced virome reads, which is within the typical range^38^. After assembly and species-level deduplication, the number of non-viral OTUs was 20 times greater than our total number of viral OTUs. vOTUs were longer and more prevalent than contaminating bacterial non-vOTUs which tended to be sample-specific. Random fragments of bacterial DNA likely became copurified along with the viral particles, explaining why they were not generally conserved between samples. Contaminant DNA species thus made up the majority of the overall sequence diversity but were shorter and less prevalent than the viruses.

*Skunaviridae*, the most abundant caudoviral family in our data, comprised only 8 complete vOTUs, and this is atypical considering the hundreds of vOTUs in most of our other abundant viral families. All reference phages belonging to the family infect *Lactococcus* while our vOTUs were predicted to infect *Streptococcus*, but this could be an artefact caused by the lack of lactococcal CRISPR spacers in our host prediction database. *Streptococcus*, although very prevalent in the children, may not be abundant enough to support the high counts of virulent *Skunaviridae*. We also did not find any strong (anti)correlation between *Streptococcus* and *Skunaviridae* counts in the data. Thus, it remains a possibility that these phages were ingested as dairy products and survived the digestive tract, as has also been proposed earlier^57^. That they end up as the most abundant viral family in our study is still surprising, but could be explained by the overall phage load in the human gut. Our epifluorescence VLP counts place gut viruses at a billion per gram of feces, or at least an order of magnitude lower than the density of gut bacteria. Our estimate is consistent with other estimates in both adults and infants^8, 58^, and such scarcity of phages in the gut, would make it all the more likely for fecal virome extractions to occasionally become flooded by ingested VLPs over proliferating ones.

In our previous study on *E. coli* phages isolated from the same samples^37^, we found that virulent coliphages were found less frequently, but were more abundant and had broader host ranges, at least at the strain level. Temperate coliphages on the contrary, were frequently isolated, but had limited host ranges. Here, we found a very similar pattern albeit on a much larger scale. Virulent phage families were more abundant but less prevalent than temperate phage families. Although we could not see a difference in host range, we did find that the temperate phage families were more genetically diverse compared to the virulent ones. This observation likely reflects frequent prophage induction from lysogenic gut bacterial strains, as shown in mouse models^59–61^, and where the induced virions do not readily infect new hosts and multiply. In viromcs, this would appear as a background of temperate phages on top of which the entry of any virulent phage, through food and water, would generate sporadic phage blooms. For our infant samples this temperate background was intense enough to overshadow the diversity of virulent phages. Possibly, in adult viromes where the GM has reached an equilibrium, the bacteria are less stressed in turn making the temperate virome less dominant. This notion is consistent with how a virulent phage core is linked to adult gut health^62^, as well as the paucity of crAssphage in infant viromes^36^.

For resolving the taxonomy of our vOTUs into genera, subfamilies, families and order-level clades we used an amino acid identity (AAI) based phylogenomic approach and applied global cutoffs after rooting. Although it has been argued that global cutoffs are not suitable for virus classification^23^ we found they worked well, and they come with the key advantage of reproducibility. The existing guideline for defining new phage families^25^ involves manual inspection of gene-sharing networks^20^, and a reproducible alternative would be preferred. One might argue that clustering phages based on the proportion of shared proteins could lead to co-clustering of phages sharing accessory rather than core genes. To this day however, phylogenomics has proven a robust method for phage classification as it resists the formidable capacity of phages to exchange genetic material.

Although the large terminase subunit (TerL) was the most conserved protein in our caudoviruses, its gene was frequently exchanged such that even members of the same viral genus would carry different TerL homologs (Figure S5). Notable examples of this phenomenon are found in *Gramvirales*. Thus, the practice^26, 28^ of using TerL phylogeny to classify caudoviral phages can sometimes produce confusing results. As shown by Yutin *et al.*^28^, *Crassvirales* is not TerL monophyletic and non-Crass phages often encode TerLs that wind up in the middle of the crAss TerL tree. The recent introduction of *ε-crassviridae* into *Crassvirales*^28^ likely illustrates this problem, as our results indicate that the family is not a crAssphage family.

Finally, we found that the latest generation of metagenome virus discovery tools such as CheckV, viralVerify and VIBRANT, in conjunction with ViromeQC should account for most sequences in one’s virome data. This recent development begins to question the continued relevance of the viral dark matter problem for DNA viromics at least. Although the sensitivities of the tools against our data never got close to 100%, most of the sequences missed by the best tools were just too short to pass the imposed quality thresholds. Thus, their predictions were good and certainly easier to obtain than manual curation. On the other hand, the performances of the alignment-free methods were very close to random with our dataset, and it appears that nucleotide-level motifs do not carry enough information to distinguish viruses from their abundant hosts.

## Conclusion

We deeply sequenced 647 infant gut viromes and tackled the viral dark matter by manual curation. The approach enabled taxonomic resolution of all viruses found, uncovering 248 viral families in total, 232 of which were novel, and most of which belong to the *Caudoviricetes* viral class. We found that temperate phages dominate the infant gut virome, while Crassphage is a minor player overshadowed by several larger novel viral orders. We used our manual data set to benchmark a series of recent virus discovery tools, and found that the dark problem is practically resolved for DNA viromes. Our comprehensive annotation of the infant gut viromes provides a framework for making biologically meaningful statistical analyses against cohort clinical phenotypes for future translational viromics research efforts.

## Acknowledgments

This work is supported by the Joint Programming Initiative ‘Healthy Diet for a Healthy Life’, specifically here, the Danish Agency for Science and Higher Education, Institut National de la Recherche Agronomique (INRA), and the Canadian Institutes of Health Research (Team grant on Intestinal Microbiomics, Institute of Nutrition, Metabolism, and Diabetes, grant number 143924). M.B.D. is recipient of graduate scholarships from the Fonds de Recherche du Québec - Nature et Technologies [259257] as well as Sentinel North and is a recipient of the Goran-Enhorning Graduate Student Research Award from the Canadian Allergy, Asthma and Immunology Foundation. JT is supported by the BRIDGE Translational Excellence Program (bridge.ku.dk) at the Faculty of Health and Medical Sciences, University of Copenhagen, funded by the Novo Nordisk Foundation (grant no. NNF18SA0034956). S.M. holds the Tier 1 Canada Research Chair in Bacteriophages [950-232136]. S.A.S is a recipient of a Novo Nordisk Foundation project grant in basic bioscience [NNF18OC0052965], along with M.A.R.

## Methods

The study was embedded in the Danish population-based COPSAC2010 prospective mother-child cohort of 736 women and their children followed from week 24 of pregnancy, with the aim of studying the mechanisms underlying chronic inflammatory diseases^63^. A total of 660 participants delivered a fecal sample 1-year after birth. Each fecal sample was mixed with 10% vol/vol glycerol broth and stored at −80°C until DNA extraction for metagenomes^31^, and virome extraction. Extraction and sequencing of virions was done using a previously described protocol^64^. Briefly, DNA from fecal filtrates enriched in viral particles was extracted and subjected to brief (30 minutes) MDA amplification and libraries were prepared following manufacturer’s procedures for the Illumina Nextera XT kit (FC-131-1096). For epifluorescence VLP estimations 10 µL of a virome sample was diluted 100-fold, fixed and deposited on a 0.02 µM filter, dried and stained with SYBR^TM^-Gold (200X), then visualised with an epifluorescence microscope using a 475 nm laser. VLPs were counted in 8 to 10 fields and multiplied over the remaining filter surface area.

Virome libraries were sequenced on the Illumina HiSeq X platform to an average depth of 3 GB per sample with paired end 2x150 bp reads. Satisfactory sequencing results were obtained for 647 samples. Virome reads were quality filtered and trimmed using Fastq Quality Trimmer/Filter (options -Q 33 -t 13 -l 32 -p 90 -q 13), and residual Illumina adapters were removed using cutadapt. Trimmed reads were dereplicated using vsearch derep_prefix and then assembled with Spades v3.10.1 using the meta flag while disabling read error correction. Decontamination clusters were generated by reducing redundancy by deduplicating the 1.5M contigs above 1 kb in size into 267k 90% ANI representatives^65^, then calling genes^66^ and aligning^67^ proteins all-against-all for building an APS tree^68^ (see Supplementary Methods). The tree was cut close to the root to obtain the decontamination clusters. Bacterial MAGs from the same samples^31^ were mined for CRISPR spacers using CRISPRDetect^69^, and the virome decontamination clusters were ranked by their extent of CRISPR targeting multiplied by sample prevalence. The protein alignment results were used to define orthologous gene clusters *de novo*^70^, and gene orthology information was used to visualise the gene contents of contigs within each decontamination cluster. The top 400 ranked clusters were inspected visually for two viral signatures, namely conservation of contig sizes and of gene content. There were diminishing returns beyond the top 400 mark and the remaining decontamination clusters were assumed to represent contaminants.

Species-level (95% ANI) deduplication of contigs into OTUs was done using BLAT^71^. Decontaminated vOTUs and reference phage species^93^ were pooled and the APS tree and gene orthology (VOGs) were recalculated. VOGs were aligned against Pfam^72^, CDD^73^, COG^74^ and TIGRFAMs^75^ using HH-suite3^76^ to gain functional annotations. The APS tree was cut using phylotreelib (https://github.com/agormp/phylotreelib) to reproduce existing phage taxonomy^28, 77^.

Family visualisations (Figure S2) were used to 1) further curate each individual vOTU in order to separate confirmable viruses that had structural VOGs, from vOTUs representing small fragments or various virus-related MGEs that did not harbour genes coding for any structural proteins. 2) The OTU length distribution within each family was plotted in a histogram with 5-kb steps to locate the right-most size peak. The 5-kb step immediately preceding this peak was set as the lower size bound for a complete or near-complete genome. 3) The family visualisations were inspected to manually remove families that were dominated by reference phages, so as to avoid interference with ongoing classification efforts. Weak families composed mainly of MGEs or fragments or having less than five vOTUs or less than two complete vOTUs were also removed. For the final version of the family visualisations available online, VOG annotations were redone against PHROGs^95^ because it was more informative.

MAG spacers, along with spacers from CRISPRopenDB^33^ and WIsH^33^ were used to generate separate host predictions for each vOTU. The three predictions were integrated using the LCA of the two most closely matching predictions, as an error-correction strategy, since all three methods would occasionally mispredict.

Bacterial contamination was estimated for each virome sample using ViromeQC^38^ along with a custom approach where we leveraged the metagenomes cognate to each virome: Reads were mapped from both fractions against 16S DNA^78^ and *cpn60*^79^ and the degree of contamination was calculated as the ratio between the two fractions. Abundances of vOTUs in each sample were determined by mapping sample reads to sample contigs using bwa mem -a^80^, then using msamtools profile to determine depth and length-normalised relative abundances with iterative redistribution of ambiguously mapped reads (https://github.com/arumugamlab/msamtools). The obtained contig abundances were then aggregated at the OTU level in order to obtain vOTU abundances per sample. vOTU abundances were aggregated at the family and order levels using phyloseq^81^ to obtain the statistics used for figures 4 and 5.

A list of VOGs matching to integrase and recombinase protein families was first curated, then used to predict whether complete vOTUs within viral families were temperate or virulent. Families where more than 95% of complete vOTUs did not harbor an integrase were deemed virulent, whereas for temperate families at least 50% of complete or incomplete vOTUs needed to encode an integrase.

The versions of virus discovery tools used for benchmarking were DeepVirFinder v1.0, VIBRANT v1.2.1, VIRSorter 1.0.6, VIRSorter2 v2.0 commit 22f6a7d, Seeker commit 9ae1488, PPR-Meta v1.1, and CheckV v.0.7.0. The random prediction was created by randomly sampling the 362k OTUs 12,500 times without replacement. The number 12,500 was chosen because it was reasonably close to our own positive set and the number of positives generated by most tools.

## Supplementary Figures

**Figure S1:**
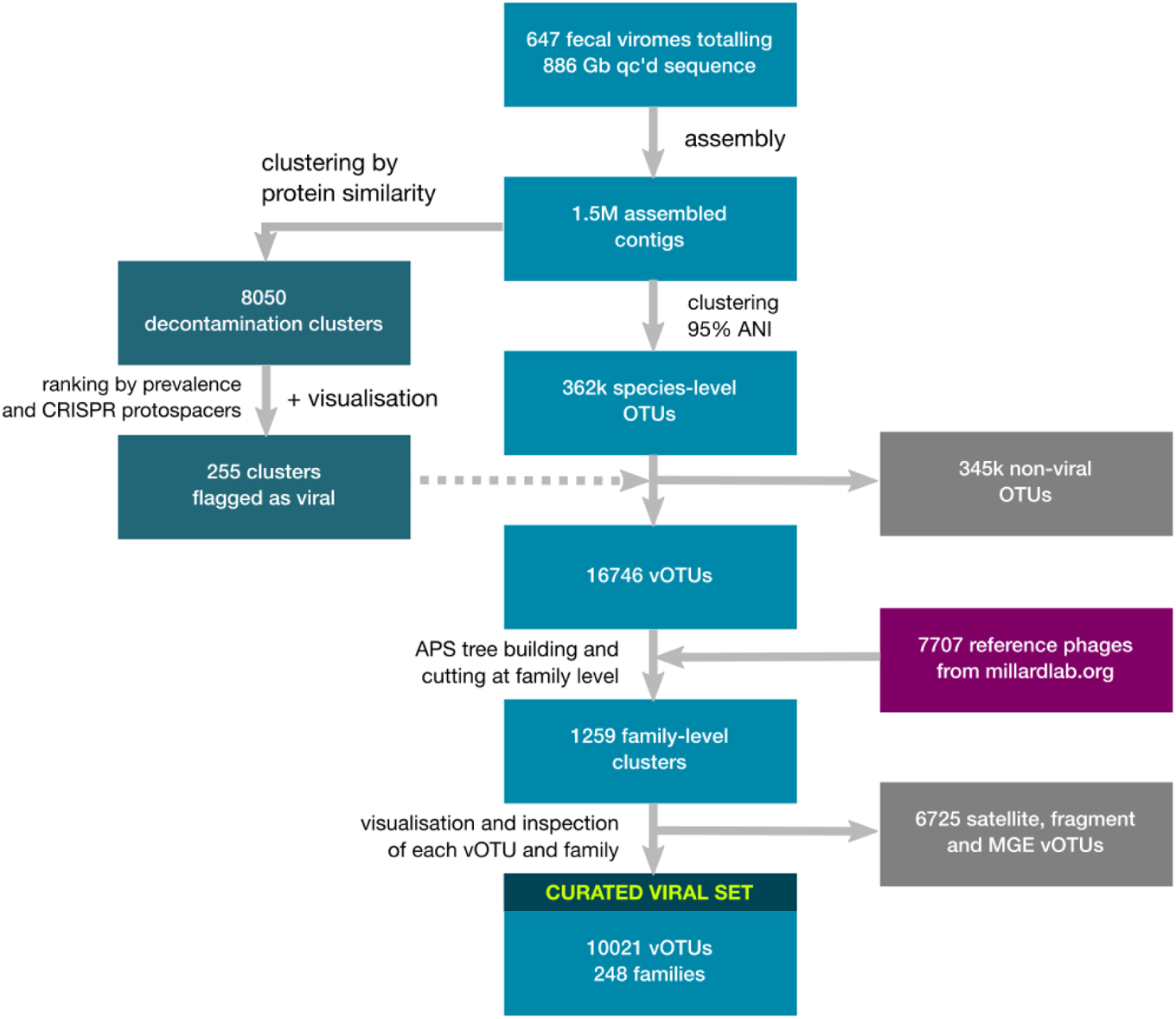
Overview of decontamination and curation procedure.

**Figure S2:**
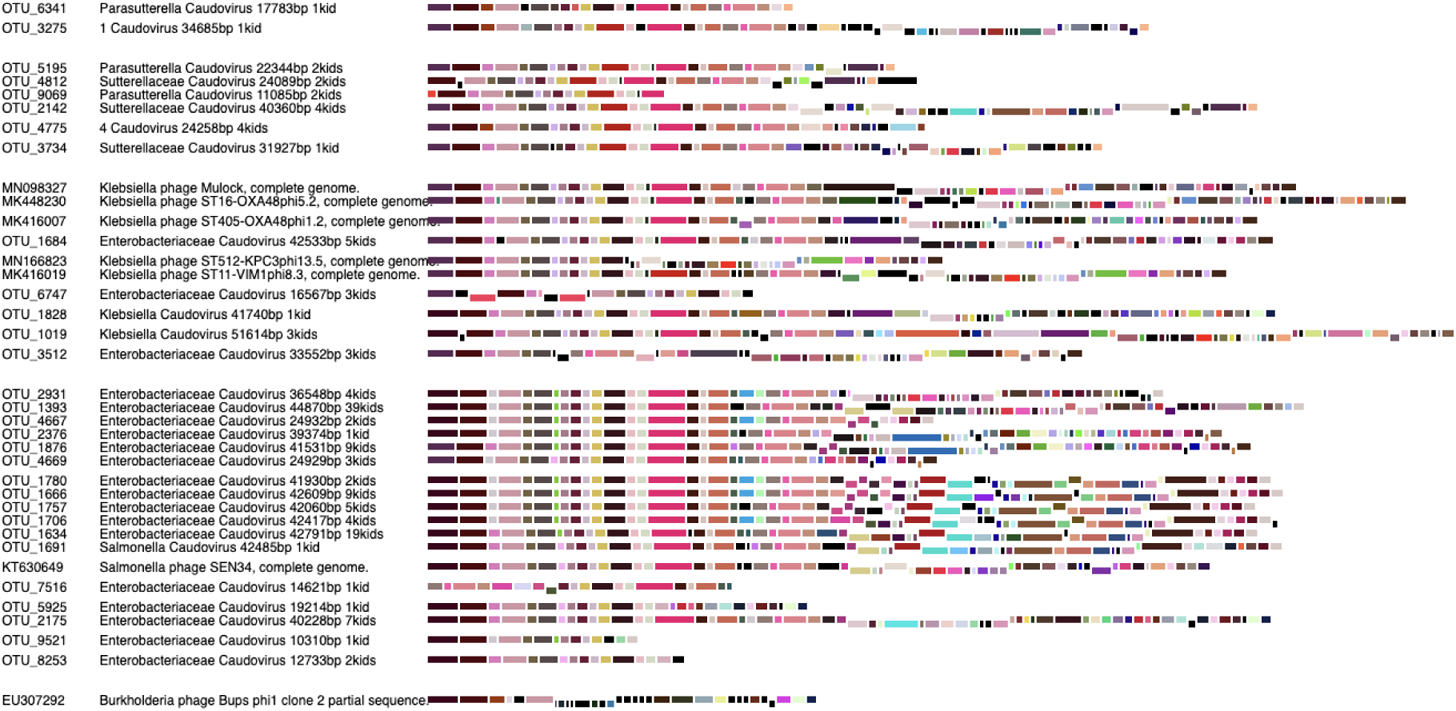
Clickable gene map of vOTUs belonging to the *Ingridviridae* family available at http://copsac.com/earlyvir/f1y/families/Ingridviridae.svg along with similar maps for the remaining 247 families, available via http://copsac.com/earlyvir/f1y/fig1.svg. Small vertical gaps between vOTUs denote genus boundaries, while large gaps denote subfamily boundaries. Ordering of the vOTUs follows the order in the APS tree and thus, related vOTUs are next to each other. ORFs are aligned vertically based on strandedness and colored by VOG affiliation. VOG definitions against the PhROGs database^95^ can be looked up by clicking on each ORF. ORF gene product (GP) numbers are displayed by mouse-over hovering. GenBank files for each vOTU can be viewed along with virus and host taxonomy by clicking on the OTU name. Caudoviral maps were inverted and zeroed according to TerL gene coordinates, while the GenBank files were not. Reference phages that belong to the same family were also included in the maps and are indicated by GenBank accession numbers.

**Figure S3:**
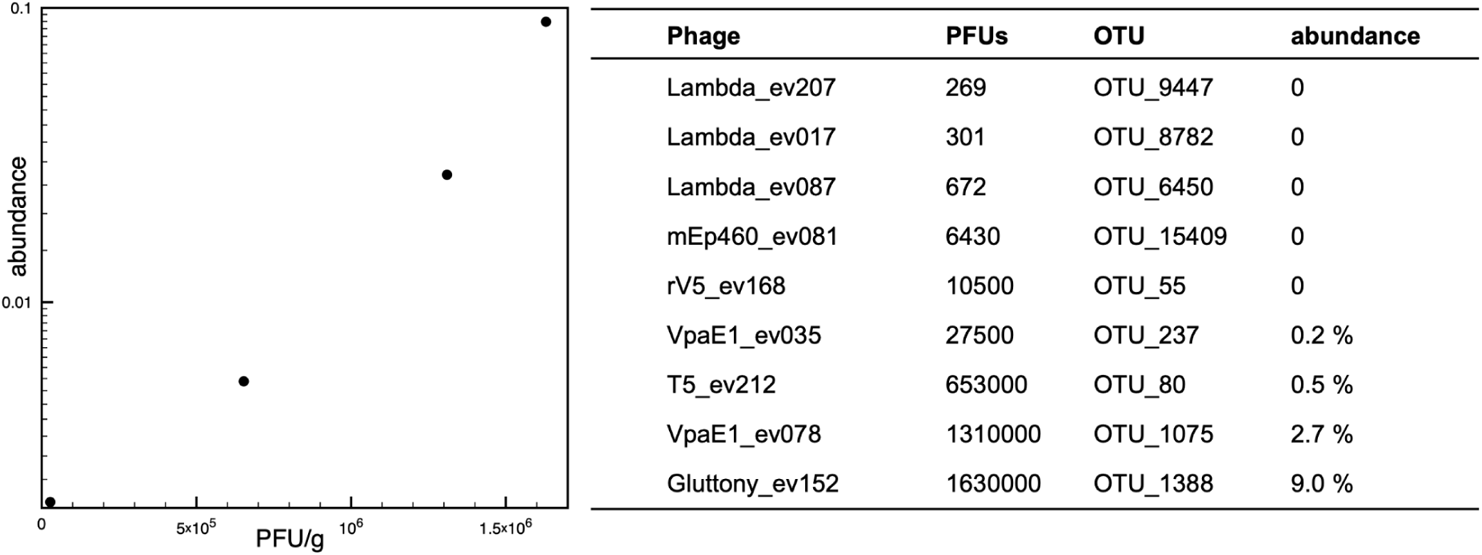
The MDA amplified viromes were quantitative for dsDNA phages. The relationship between experimentally determined PFU/g of feces for isolated coliphages^37^ and virome abundances for their corresponding vOTUs in the same samples is shown. The agreement between the two estimates supports that the dsDNA virome sequencing was quantitative despite its insensitivity to rare phages.

**Figure S4:**
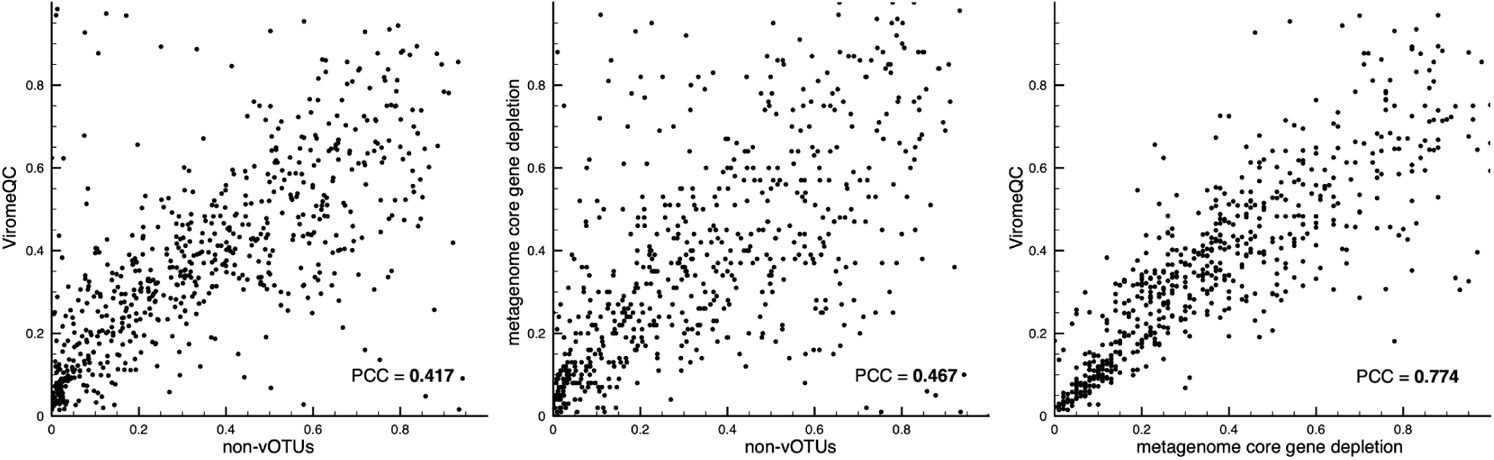
Comparison of three approaches for estimating the proportion of bacterial contamination. Each graph has 647 dots, one for each sample. Axes denote the proportion of bacterial contamination as estimated by the indicated method. Each graph is a pairwise comparison of two different methods. A) non-vOTU mappings vs. ViromeQC B) non-vOTU mappings vs. metagenome core gene depletion C) metagenome core gene depletion vs. ViromeQC. Pearson correlation coefficients are given for all three comparisons.

**Figure S5:**
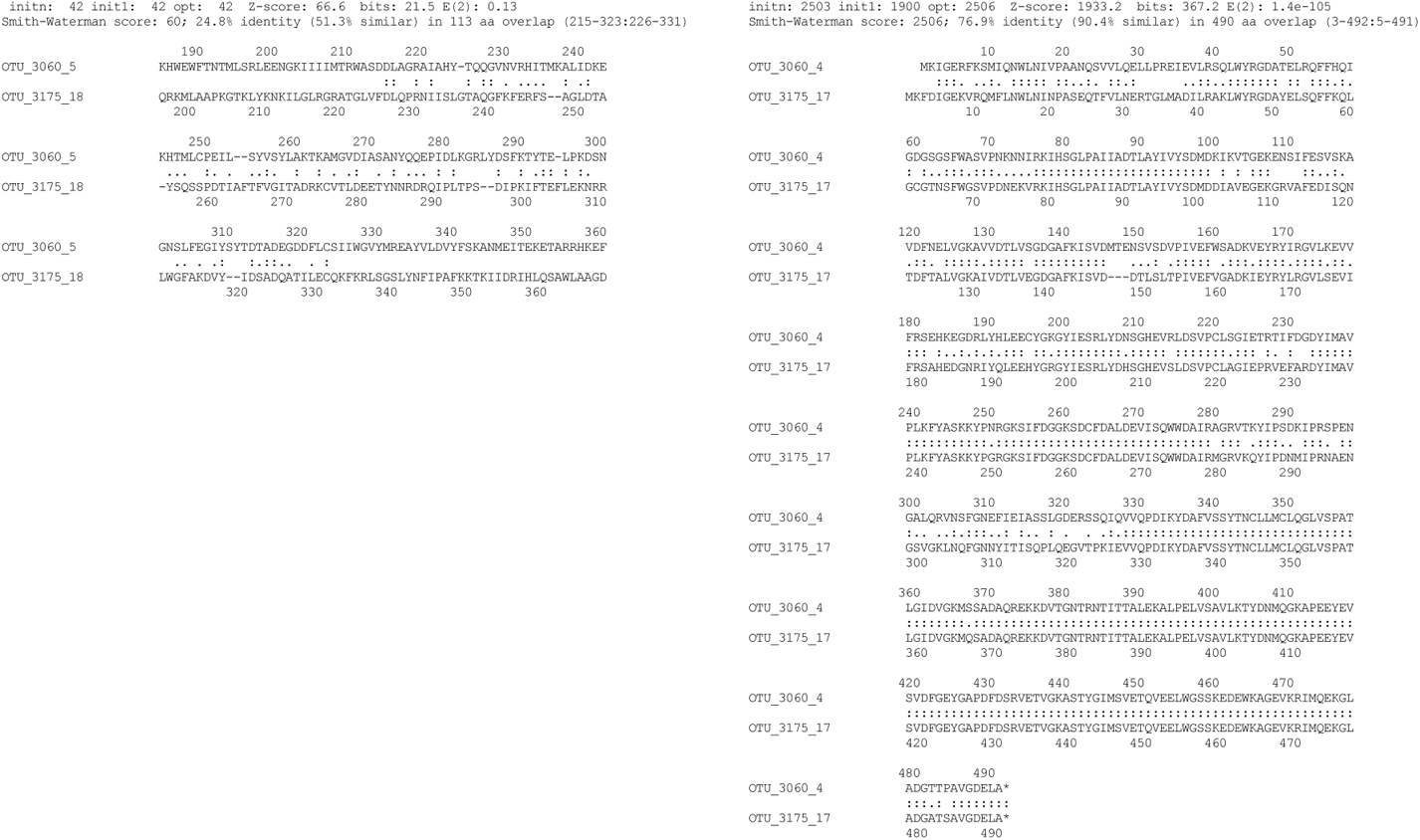
Alignments of TerL and the portal protein for two viruses belonging to the same genus within the *Jeppeviridae* family. The 75% amino acid identity for the portal protein alignment is within the expected range for phages of the same genus, whereas the terminase proteins are so disparate that gene exchange is the most likely explanation.

## Supplementary Methods

### The COPSAC2010 birth cohort

The study was embedded in the Danish population-based COPSAC2010 prospective mother-child cohort of 736 women and their children followed from week 24 of pregnancy, with the primary clinical endpoint persistent wheeze or asthma^30^.

### Fecal sample collection, transport and storage

Of the 700 children part of the COPSAC2010 cohort, 660 delivered a fecal sample 1-year after birth. Fecal samples were collected either at the research clinic or by parents at home using detailed instructions. Upon arrival, each fecal sample was mixed with 10% vol/vol glycerol broth and stored at −80°C until virome extraction.

### Virome extraction and library preparation

Preparation of fecal samples, and extraction and sequencing of virions was done using a previously described protocol^64^. Briefly, viral-associated DNA was subjected to brief MDA amplification and libraries prepared following manufacturer’s procedures for the Illumina Nextera XT kit (FC-131-1096). Libraries were sequenced, paired-end, on the Illumina HiSeq X platform.

### VLP counts by epifluorescence

A volume of 10 µL per virome sample was diluted 100-fold in SM buffer, fixed with 0.5% glutaraldehyde, and frozen in liquid nitrogen. The sample was thawed, deposited on a 0.02 µM pore size membrane (Anodisc 25, Anopore, Whatman) using a filtering device. Next, the filter was dried, incubated on a 70 µL drop of SYBR^TM^-Gold (200X) in the dark for 15 minutes, dried, and mounted on a microscope slide, together with fluoromount and antifade. VLP dots were visualized using an epifluorescence microscope equipped with an ORCA camera, and a 475 nm excitation laser. Eight to ten fields (1344x1024 pixels) were imaged and fluorescent dots counted. VLP counts on the total filter surface were deduced by multiplying the average field count by the number of fields over the total membrane surface (60 493).

### Virome assembly and gene calling

Virome libraries were sequenced to an average depth of 3 GB per sample with paired end 2x150 bp reads. Satisfactory sequencing results were obtained for 647 samples. Virome reads were first quality filtered and trimmed using Fastq Quality Trimmer/Filter (options -Q 33 -t 13 -l 32 -p 90 -q 13), and residual Illumina adapters were subsequently removed using cutadapt. The resulting trimmed reads were dereplicated using vsearch derep_prefix and then assembled with Spades v3.10.1 using the meta flag and disabling read error correction. In order to reduce sequence redundancy for the subsequent decontamination step, all 1.5M contigs above 1 kb in length were clustered at 90% ANI and coverage using a previously described approach^65^. Longest representatives from each of the 267k clusters were subjected to gene calling^66^ and the resulting protein sequences were used to build an APS tree^68^ and for *de novo* VOG definition^70^ (more details below).

### The aggregate protein similarity (APS) tree

Sharing features with DICE^82^ and VIPtree^83^, the APS tree methodology was first developed^68^ for systematic and automated classification of prokaryotic defence systems and mobile genetic elements that were traditionally classified by manual inspection. The tree was found to reproduce existing manual classes and sub-classes when cutting it with global cutoffs^68, 84^. The tree also predicted deeper relationships between classes that were experimentally validated later^85^. The method uses as input, an all-against-all protein alignment search, preferably with sensitive alignment software like FASTA^67^, although BLASTP or DIAMOND also work. For all viruses being compared, all constituent viral proteins are searched against each other. Next, virus to virus similarity scores are tallied by aggregating the alignment scores for all their constituent proteins. Next, using the Bray-Curtis dissimilarity^86^, a distance matrix is constructed between all viruses, which is then used as input for constructing a neighbour joining tree. We used RapidNJ (https://github.com/somme89/rapidNJ) for the latter. After manually rooting the tree, it can be cut at fixed distances from the root to reproduce classes at required levels. To this end we developed “treetool” described further below. Using more sensitive alignment software (like FASTA) generates more reliable distant protein alignments, making deeper cuts less noisy. In this study we found that cuts down to the viral order-level were reproducing existing and proposed taxonomy, making such cuts useful for defining novel taxa as well.

### Protein annotation, clustering and visualisation

Protein coding genes for representative contigs during initial decontamination, and for vOTUs and reference phages later were predicted using Prodigal^66^. All protein sequences were subject to an all-against-all sequence alignment using FASTA^67^. Protein alignments between viruses were used to cluster viruses into taxa using the APS tree as described above and below. But the alignments were also used to define VOGs *de novo* using a previously described orthology detection pipeline involving protein alignment coverage cutoffs^70^ and Markov Clustering^87^. Multiple sequence alignments (MSAs) and phylogenetic trees were constructed^88^ from the protein sequences corresponding to each VOG. MSAs were used for profile-profile alignments against the Pfam^72^, COG^74^, CDD^73^, TIGRFAMs^75^ and PhROGs^95^ databases using HHblits^76^ in order to annotate them and determine their functions. Gene contents of the contigs were visualised along with other contigs belonging to the same viral clusters by coloring genes by orthology and hyperlinking profile-profile results in SVG graphics as in Figure S2. The SVG files were generated using a custom bash script. The graphics were browsed manually during the manual decontamination, species curation and family curation steps (Figure S1).

### Manual decontamination of assembled contigs

All proteins from the 267k 90% ANI contig cluster representatives were subject to an all-against-all FASTA^67^ search that was used for defining VOGs^70^. The search results were also used to construct an APS tree that was cut close to the root in order to yield 8050 decontamination clusters of contig representatives. SVG graphics were generated for each decontamination cluster by visualising each individual representative within it. This was done by using the gene orthology information generated earlier to color rectangles corresponding to encoded genes. The decontamination clusters were then ranked by their sample prevalence and extent of CRISPR targeting from our MAGs^31^. SVG graphics for the first 400 ranked decontamination clusters were manually inspected and the 255 clusters resembling viral families were flagged. Viral clusters were recognised by having conserved genome sizes with the majority of the genes and synteny being largely conserved between most members within the cluster. Plasmid clusters, on the other hand, were often heterogeneous in size and extremely variable with regard to gene content, and would encode mobilisation proteins, relaxases or Type IV secretion systems. Bacterial contaminant clusters were made up of short contigs sharing one or two genes, but otherwise variable in size with any remaining genes being disparate. Clusters ranking lower than the top 400 were not inspected further due to diminishing returns, as most additionally inspected clusters at this point resembled bacterial contamination. For the 255 flagged viral clusters, contigs smaller than 10 kb were discarded unless the apparent complete genome size for that cluster was shorter, as for the ssDNA viruses.

### Species-level OTU delineation

Species-level deduplication was necessary because very similar viruses from different samples were assembled into separate contigs, as assemblies were carried out separately per sample. These had to be merged into single OTUs and a representative for each OTU had to be chosen. This was done by comparing all assembled contigs from all samples to each other, to find clusters of very similar contigs. In order to account for incomplete assemblies we selected the longest version as the OTU representative sequence. However, the longest version of a species was sometimes a chimeric assembly, where two closely related viruses had been merged into a long contig e.g. containing two copies of each gene. To avoid selection of chimeras as OTU representatives, all contigs with more than 110% self-similarity (judging from self-alignment score over length) were flagged as potentially repetitive and were not selected as a representative. In such cases the next-longest sequence was selected to represent the OTU. A variety of tools were tested for comparing the contigs and generating OTUs at the 95% species-level ANI, including cd-hit-est^89^, BB dedupe^90^ and nucmer^91^. The most consistent results were obtained using BLAT^71^ and setting a cutoff on the alignment score. This was done by requiring a score of 90% of what the shorter sequence obtained against itself, while aligned against a longer sequence, in which case the two were merged. The 90% score would correspond on average to 95% identity at 95% coverage, but at extremes in either direction could also occasionally include 100% identity at 90% coverage or 90% identity at 100% coverage. This flexibility was a compromise offset by the high accuracy of BLAT vs. other tools tested. The clustering step itself was carried out with a perl one-liner applied on top of an all-against-all BLAT output, where all assembly contigs had been used as input.

### PhyloTreeLib.py for exploring and cutting the APS tree

phylotreelib (https://github.com/agormp/phylotreelib) is a Python library developed in-house for manipulating or extracting information from phylogenetic trees. Here, we used phylotreelib.py via its command line front-end treetool.py (https://github.com/agormp/treetool). After constructing the APS tree it was visually inspected in FigTree (https://github.com/rambaut/figtree/) to find a suitable outgroup. Suitable outgroups comprised branches originating directly from the stem of the tree. The stem was easily visible due to the high diversity of sequences used to construct the APS tree. Next, treetool.py was used with the --root option to root the APS tree based on the selected outgroup. Next, a list of accession numbers was made containing all members of the *Herelleviridae* family, along with separate lists for members of different subfamilies and genera within the *Herelleviridae*. Another list was made containing all known crAssphages at the time of the analysis. Then, treetool.py was used along with the --cladeinfo option to retrieve the distances from the root to the branch encompassing the leaves in the lists. Next, treetool.py’s --clustcut option was used to cut the tree at the above distances in order to obtain clades of vOTUs and reference phages corresponding to viral orders, families, subfamilies and genera. The distances we used to cut the tree were 0.250, 0.125, 0.04 and 0.025 respectively for genus, subfamily, family and order-level clades. Thus the approximate minimum AAI and coverage required for two viruses to belong to the same clade could be derived from those cut distances to be on average 70%, 50%, 28% and 22%.

### Generation of families and species-level curation

To find viral species corresponding to the flagged viral decontamination clusters, all 1.5M contigs were clustered at 95% ANI, resulting in 362k OTUs. The 8327 viral sequences from the decontamination step were aligned against 95% ANI vOTUs requiring a 50% BLAT score (∼75% ANI at ∼75% coverage). The 16,746 95% ANI vOTUs recruited in this manner were pooled with 95% ANI dereplicated reference phages from millardlab.org^93^. Protein coding genes were annotated for all sequences and subject to an all-against-all FASTA search. The *de novo* VOGs and APS tree were recomputed. The tree was rooted manually using phylotreelib, and the *Herelleviridae* family-level root distance was measured using treetool’s cladeInfo function. The tree was cut at that distance yielding 1259 viral family-level clades. In order to make sure that the large number of families was not a result of accidental recruitment of additional contaminant contigs during the 75% ANI matching step from above, each vOTU and reference phage within each family-level clade was visualised using the VOG information as seen in Figure S2. Next, each individual vOTU was manually flagged as belonging to either one of the categories virus, putative satellite, putative MGE or unknown contaminant using the following guidelines. To be considered viral, a vOTU needed to encode at least one structural protein along with an additional viral protein. vOTUs that did not live up to these criteria were considered MGEs if they encoded at least one protein indicative of a mobile lifestyle (e.g. integrases or replication proteins), or if they were conserved in size and gene content across multiple samples. The rationale for the latter was that random segments of contaminating DNA should not be conserved in size and gene content, while any mobile DNA would be expected to do so. Unknown viruses that encoded novel structural proteins could thus end up in the MGE category. MGEs that co-clustered within viral family-level clades, or contained at least two non-structural viral proteins were classed as putative satellites. vOTUs that did not live up to any criteria were classed as unknown contaminants. A class of non-coding sequence was particularly prevalent in the unknown group and were flagged separately.

### Final family curation

All 1259 family-level clades were manually inspected to make sure that they looked like true families instead of being artefacts of the methods employed. The majority of the family-level clades consisted of singleton vOTUs or reference phages, and these were discarded. Additionally, most families consisting mostly of reference phages were discarded to avoid interference with independent classification efforts. Finally, families that looked unconvincing because they consisted of mostly MGEs, satellites, or genome fragments were also removed, leaving 248 curated families in the end. For each curated family, a histogram of vOTU length was made using a bin width of 5 kb. The minimum genome completion size for each family was determined by inspecting each histogram and choosing the 5-kb bin immediately preceding the rightmost peak. This size cutoff was used subsequently to select all complete and near-complete vOTUs within each family. Families that did not have enough vOTUs to yield a convincing genome size peak were flagged as having an unknown completion size.

### Host prediction

As part of a separate effort^31^, metagenomes from the same samples had been assembled and binned into MAGs that had taxonomies predicted using GTDB-tk. CRISPR arrays were predicted on each MAG using CRISPRdetect^69^, and the spacers were pooled and used as a host prediction database, alongside the CRISPR spacer database^33^. The host prediction algorithm from the CRISPR spacer database was used to predict hosts for each vOTU using either spacer database. An additional host prediction was made using WIsH v1.0^34^, against a database of complete bacterial genomes with null-models calibrated against all reference phage sequences available at millardlab.org^91^. The best maximum-likelihood WIsH prediction per vOTU having a P-value under 0.05 was used. Host predictions from all three methods were translated into NCBI taxonomy identifiers and integrated by taking the LCA of the two closest genus-level predictions. If WIsH predicted *Bacteroides*, public spacers *Clostridium* and metagenome spacers *Faecalibacterium*, then the host was set to order clostridiales, being the LCA of the two most closely matching predictions. The rationale for this approach was to absorb occasional false positives stemming from either of the three methods, as all three methods had been seen to make false predictions on separate occasions. However, most predictions were unanimous at the genus-level.

### Quantification of bacterial contamination

Two approaches were used for quantifying the amount of bacterial contamination in the virome samples. In the first approach, ViromeQC was run on the QC’d virome reads, and the proportion of bacterial contamination per sample was determined by taking the inverse of the “virus enrichment ratio” as reported by ViromeQC.

In the other approach, we wanted to take advantage of our coupled metagenomes for each virome sample, hoping to gain a more accurate estimate than ViromeQC, which does not utilise such additional information. We did this by first searching reads from the virome with the Rfam 16S rDNA model (RF00177). Then the same was done for reads from cognate metagenomes. The ratio between the proportion of reads mapping to RF00177 in the virome over the same proportion for the metagenome was counted as the estimated bacterial contamination ratio by 16S content. In parallel, reads from both the viromes and metagenomes were mapped against a database of the bacterial *cpn60* core gene from cpnDB^79^, using bowtie2^92^. As we did for the 16S-based contamination estimate, the proportion of reads mapping to this database in the virome over the metagenome for each sample was counted as the estimated bacterial contamination ratio by *cpn60* content. Although the two estimates, by 16S and *cpn60*, were in general agreement, they also sometimes differed. The mean between the two estimates was used as the “metagenome core gene depletion” contamination-estimate used in Figure S4.

### Estimating sample vOTU abundances

vOTU counts were generated by mapping the reads of each sample against only the assembled contigs from that particular sample. Mapping was performed using the `bwa mem -a` flag^80^. Contig counts were generated with `msamtools profile` (https://github.com/arumugamlab/msamtools), which computes relative abundances of each contig, normalising by contig length and mapping depth while iteratively redistributing non-uniquely mapping reads. Abundances of contigs corresponding to vOTUs were aggregated by leveraging the 95% ANI dereplication cluster membership information computed previously. Counts for contigs that did not correspond to any vOTU were assumed to represent contaminant DNA and gathered as an independent contamination estimate. The vOTU to sample relative abundance matrix, or OTU table, was loaded into a phyloseq object in R and agglomerated at the viral family and order-levels to compute the statistics used in Figures 4 and 5.

### Prediction tool benchmarking

The versions of virus prediction tools used were DeepVirFinder v1.0, VIBRANT v1.2.1, VIRSorter 1.0.6, VIRSorter2 commit 22f6a7d, Seeker commit 9ae1488, PPR-Meta v1.1, and CheckV v.0.7.0. Some of these tools were designed for virus discovery in metagenomes, while others meant for virome decontamination, but the two are related given the high amount of bacterial contamination in most viromes published to date^38^. Despite our own efforts to obtain high-quality virome samples, the vast majority of our OTUs were bacterial contamination rather than viral. For this reason no distinction was made between the two types of tools. For the tools that required cutoffs, we used typical cutoffs used by the research community, or if such information was not available, we tried to set a cutoff that would generate a number of positive predictions that reasonably matched our own positive set. Since our data set is so skewed, with 10,021 positive viral sequences versus over thirty times more negatives, performance measures can be biased, which is why we included the “kappa” metric, designed to address this issue, as well as a random prediction for comparison. The random prediction was created by randomly subsampling the 362k OTUs 12,500 times without replacement. The number 12,500 was chosen because it was reasonably close to our own positive set and the number of positives generated by most tools.

## Notes

http://copsac.com/earlyvir/f1y/fig1.svg

http://copsac.com/earlyvir/f1y/taxtable.html

http://copsac.com/earlyvir/f1y/vogs.html

http://copsac.com/earlyvir/f1y/families/Ingridviridae.svg

### Competing Interest Statement

The authors have declared no competing interest.

